# New mutant mouse models clarify the role of NAIPs, phosphorylation, NLRP3, and tumors in NLRC4 inflammasome activation

**DOI:** 10.1101/765313

**Authors:** Jeannette L. Tenthorey, Roberto A. Chavez, Thornton W. Thompson, Katherine A. Deets, Russell E. Vance, Isabella Rauch

## Abstract

The NAIP/NLRC4 inflammasome is a cytosolic sensor of bacteria that activates Caspase-1 and initiates potent downstream immune responses. Structural, biochemical, and genetic data all demonstrate that the NAIP proteins act as receptors for specific bacterial ligands, while NLRC4 is a downstream adaptor protein that multimerizes with NAIPs to form a macromolecular structure called an inflammasome. However, several aspects of NLRC4 biology remain unresolved. For example, in addition to its clear function in responding to bacteria, NLRC4 has also been proposed to initiate anti-tumor responses, though the underlying mechanism is unknown. NLRC4 has also been shown to be phosphorylated on serine 533, and this modification was suggested to be important for NLRC4 function. In the absence of S533 phosphorylation, it was further proposed that another inflammasome component, NLRP3, can induce NLRC4 activation. We generated a new *Nlrc4*-deficient mouse line as well as mice encoding phosphomimetic S533D and non-phosphorylatable S533A NLRC4 proteins. Using these genetic models *in vivo* and *in vitro*, we fail to observe a role for phosphorylation in NLRC4 inflammasome function. Furthermore, we find no role for NLRP3 in NLRC4 function, or for NLRC4 in a model of melanoma. These results simplify and clarify our understanding of the mechanism of NAIP/NLRC4 activation and its biological functions.

## INTRODUCTION

Inflammasomes are innate immune multi-protein complexes that assemble to activate pro-inflammatory caspases following the detection of cytosolic pathogen-or stress-associated stimuli (Martinon et al., 2002; Lamkanfi and Dixit, 2014; Rathinam and Fitzgerald, 2016). The NAIP/NLRC4 inflammasome is one of the best characterized inflammasomes. NLRC4 was identified in 2001 as a CARD (Caspase Activation and Recruitment Domain)-containing protein that can activate Caspase-1 (Poyet et al., 2001). NLRC4 was found to activate Caspase-1 after cytosolic invasion by flagellated or type 3 secretion system (T3SS)-expressing bacterial pathogens (Franchi et al., 2006; Miao et al., 2006, 2010; Mariathasan et al., 2004; Zamboni et al., 2006). Subsequent genetic, biochemical, and structural studies established that activation of NLRC4 requires NAIP proteins, which act as sensors to detect bacterial flagellin or T3SS needle or rod proteins (Kofoed and Vance, 2011; Zhao et al., 2011; Rauch et al., 2016; Zhao et al., 2016; Tenthorey et al., 2017). Binding of bacterial ligands by NAIPs leads to structural changes that allow for the recruitment and oligomerization of NLRC4, which acts as a downstream adaptor to recruit and activate the Caspase-1 protease (Hu et al., 2015; Zhang et al., 2015; Tenthorey et al., 2017). Activated Caspase-1 cleaves the cytokines IL-1β and IL-18 as well as a pore-forming protein called Gasdermin D, and this cleavage is required for maturation (Rathinam and Fitzgerald, 2016; Lamkanfi and Dixit, 2014; Shi et al., 2015; Kayagaki et al., 2015). Mature Gasdermin D causes pyroptotic host cell death to release active IL-1β and IL-18, leading to potent inflammatory responses to control infection.

Despite nearly two decades of study, several aspects of NLRC4 activation and function remain unclear. First, NLRC4 may have biological roles outside of pathogen sensing, such as in immune sensing or control of tumors. Several inflammasomes, including NLRP3, NLRP1, and AIM2, have been reported to largely protect against tumor growth (Karki et al., 2017). Recent reports have also implicated NLRC4 in tumor progression. NLRC4 and Caspase-1 were found to promote the metastasis of breast and colon cancer in obese mice, but not those fed on normal-fat diets (Kolb et al., 2016; Ohashi et al., 2019). In contrast, NLRC4 and Caspase-1 were reported to protect mice from chemically-induced colon cancer (AOM/DSS) (Hu et al., 2010), although another report found that Caspase-1 but not NLRC4 was protective in the same model (Allen et al., 2010). Intriguingly, a recent paper found that *Nlrc4*^*−/−*^ mice but not *Caspase1*^*−/−*^ mice are susceptible to implanted B16 melanomas (Janowski et al., 2016), suggesting that NLRC4 might mediate tumor control independently of its prototypical Caspase-1 pathway. At least some of the differences in these results might derive from the use of varying tumor cell lines or induction models. However, it is currently unclear what might be driving the activation of NLRC4 in tumors or tumor-associated macrophages, as the only known agonists of NLRC4 are cytosolic bacterial proteins.

An additional unresolved question is the role of phosphorylation in NLRC4 inflammasome activity. Phosphorylation of NLRP3 by JNK1 is essential for NLRP3 inflammasome assembly (Song et al., 2017), providing precedent for a phosphorylation-based checkpoint on inflammasome signaling. NLRC4 is also phosphorylated, at serine 533 (S533), during *Salmonella* infection, and *Nlrc4*^*−/−*^ macrophages reconstituted with non-phosphorylatable NLRC4 (alanine mutant S533A) failed to induce an inflammasome response to *Salmonella* infection (Qu et al., 2012). Conversely, it was reported to be difficult to reconstitute *Nlrc4*^*−/−*^ macrophages with a phosphomimetic NLRC4 (aspartic acid mutant S533D), suggesting that NLRC4 S533D might be constitutively active for induction of pyroptosis in the absence of infection. Thus, NLRC4 phosphorylation was proposed to be both necessary and sufficient for NLRC4 inflammasome activation. The authors further identified protein kinase C delta (PKCδ) as required for NLRC4 phosphorylation and activation (Qu et al., 2012). The same group subsequently generated mice with a homozygous S533A mutation in the endogenous *Nlrc4* gene (Qu et al., 2016). In contrast to the prior claim that phosphorylation was critical for NLRC4 activation, macrophages from knock-in mice expressing NLRC4-S533A exhibited only a partial defect in NLRC4 activation after *Salmonella* infection.

However, the residual inflammasome signaling in NLRC4-S533A cells was eliminated by deletion of the gene encoding NLRP3 (Qu et al., 2016), another inflammasome that participates in *Salmonella* sensing (Broz et al., 2010). The authors proposed that NLRP3 is recruited to NLRC4 to mediate inflammasome activation by the S533A mutant. However, it is currently unclear why phosphorylation might be required for optimal NLRC4 signaling, as S533 is not near the NLRC4 oligomerization or Caspase-1-recruitment interfaces (Tenthorey et al., 2017; Hu et al., 2015; Zhang et al., 2015). Nor is it clear how NLRP3 is recruited and contributes to NLRC4 signaling.

Critically, the above studies did not address whether S533 phosphorylation acts in concert with or independently of NAIPs to mediate NLRC4 activation. The *Salmonella* T3SS was required for NLRC4 phosphorylation (Qu et al., 2012). In addition, transfected flagellin was sufficient to induce NLRC4 phosphorylation (Qu et al., 2016). Since NAIP proteins are required for flagellin and T3SS proteins to activate the NLRC4 inflammasome (Kofoed and Vance, 2011; Zhao et al., 2011; Lightfield et al., 2008; Zhao et al., 2016; Rauch et al., 2016), the simplest model to explain the above results would be a linear pathway in which NAIP activation leads to NLRC4 phosphorylation and activation. However, NLRC4 was reported to be phosphorylated in flagellin-transfected *Naip5*^*−/−*^ cells, implying the existence of a NAIP-independent pathway for NLRC4 phosphorylation (Matusiak et al., 2015). The lack of requirement for NAIP5 in flagellin-triggered NLRC4 phosphorylation could be due to redundancy between NAIP5 and NAIP6 for detection of flagellin (Kofoed and Vance, 2011). Regardless, cells lacking NAIP5 did not activate Caspase-1 in response to flagellin stimulation, despite NLRC4 phosphorylation (Matusiak et al., 2015). Furthermore, NLRC4 was phosphorylated in an inactive, monomeric crystal structure of NLRC4 (Hu et al., 2013). These results imply that S533 phosphorylation is not sufficient to mediate NLRC4 activation, in contrast to the prior suggestion (Qu et al., 2016) that a phosphomimetic S533D NLRC4 mutant is constitutively active. In addition, the proposal that PKCδ mediates NLRC4 phosphorylation has also been questioned, since cells from *Prkcd*^*−/−*^ mice lacking PKCδ exhibit normal or even enhanced inflammasome activity (Suzuki et al., 2014).

In an effort to clarify the role of S533 phosphorylation in NLRC4 activation, we used CRISPR/Cas9 to generate three new lines of NLRC4 mutant mice: a line of NLRC4 null mice (*Nlrc4*^*−/−*^), a line of mice harboring a serine 533 phosphomimetic (S533D) mutation (*Nlrc4*^*S533D/S533D*^), and a line of mice with non-phosphorylatable (S533A) NLRC4 (*Nlrc4*^*S533A/S533A*^). These mouse lines were all generated on a pure C57BL6/J background, in contrast to another widely used *Nlrc4*^*−/−*^ mouse line which was generated on a C57BL6/N background (Mariathasan et al., 2004). Comparing our new *Nlrc4* mutant mice to C57BL6/J wildtype controls, we could not observe a major role for NLRC4 phosphorylation or NLRP3 in NLRC4 activation. In addition, we found no role for NLRC4 in control of B16 melanoma progression. Our results help to clarify the function and mechanism of activation of NLRC4 and establish NAIP proteins as the only known upstream activators of NLRC4.

## RESULTS AND DISCUSSION

To study the role of phosphorylation in NLRC4 activation, we generated multiple mutant mice via CRISPR/Cas9. We used a single guide RNA designed to induce Cas9 cleavage close to the codon for serine 533 (Fig. 1). The guide was injected into fertilized C57BL/6J oocytes together with Cas9 mRNA and two single-stranded DNA oligonucleotides as templates for homology directed repair. These oligonucleotides contained either the S533A or S533D mutation, as well as several silent mutations to avoid re-cutting of the successfully repaired allele. Founder mice were screened for the presence of the desired alleles as well as frame shifts leading to a stop codon. To separate alleles and create individual lines, and to remove potential off-target mutations, founders were backcrossed at least 3 times to C57BL/6J mice before crossing heterozygotes to generate homozygous mutant mice. Thus, we generated three new mouse lines: (1) a mouse line with a 1 bp insertion in codon 529 that leads to a premature stop at codon 542 (Fig. 1A, official name *Nlrc4*^*em1Vnce*^, in this paper *Nlrc4*^*−/−*^); (2) a mouse line harboring a homozygous non-phosphorylatable S533A mutation (Fig. 1B, official name *Nlrc4*^*em2(S533A)Vnce*^, in this paper *Nlrc4*^*S533A/S533A*^); and (3) a mouse line harboring a homozygous phosphomimetic S533D mutation (Fig. 1C, official name *Nlrc4*^*em3(S533D)Vnce*^, in this paper *Nlrc4*^*S533D/S533D*^). All mouse lines were viable and fertile, and we did not observe any obvious abnormalities.

**Figure 1.**
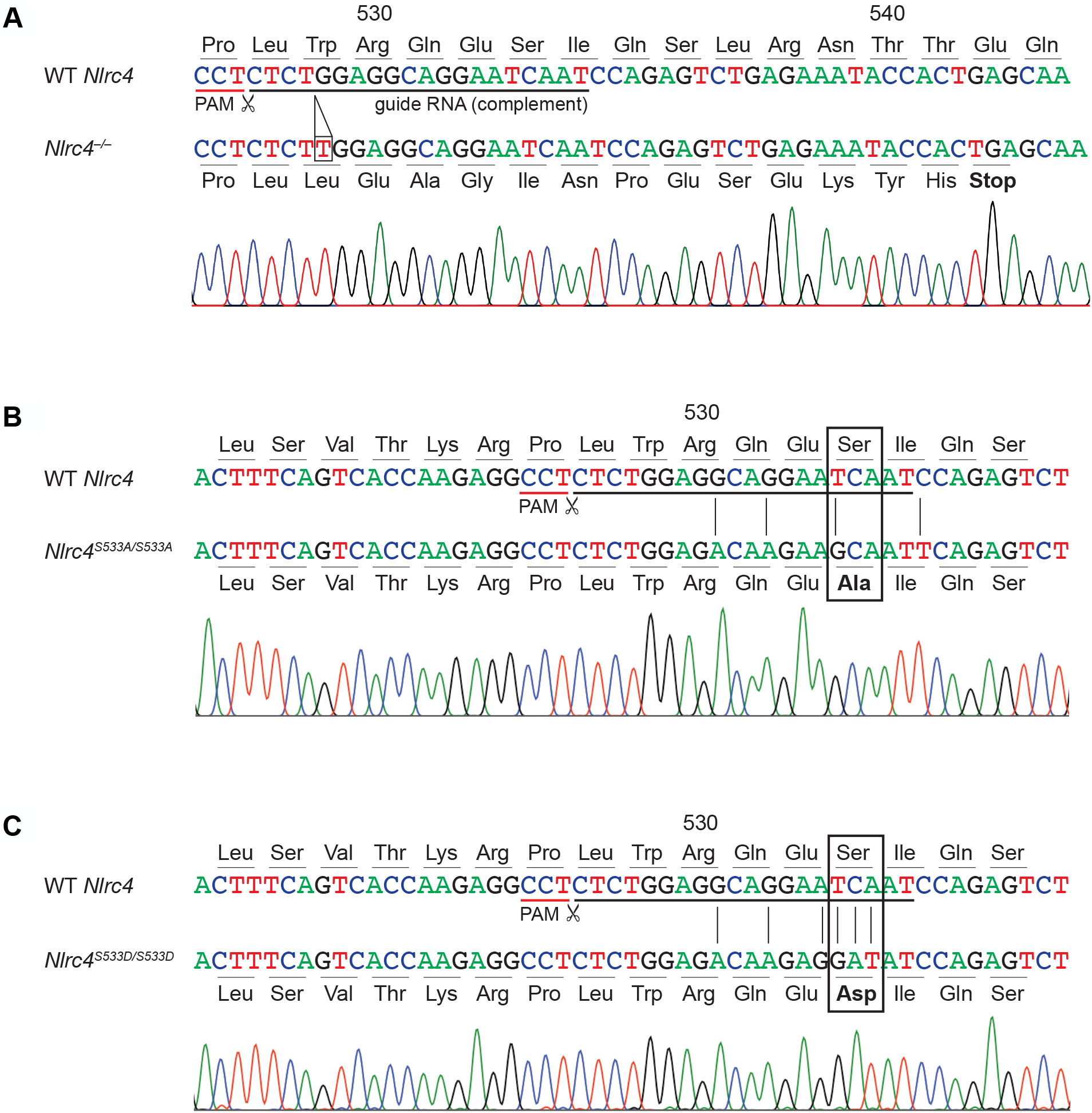
CRISPR/Cas9 targeting of *Nlrc4*. The sequence of the *Nlrc4* locus in WT C57BL6/J is compared to CRISPR/Cas9-generated (A) *Nlrc4*^*−/−*^, (B) *Nlrc4*^*S533A/S533A*^, and (C) *Nlrc4*^*S533D/S533D*^ mice. Sanger sequences traces are shown for mutant mice. The guide RNA sequence and protospacer-adjacent motif (PAM) for CRISPR/Cas9 targeting, as well as non-silent base exchanges, are indicated.

To confirm the successful inactivation of *Nlrc4* and to test the functionality of the non-phosphorylatable and the phosphomimetic alleles, we generated bone marrow-derived macrophages from these mouse lines. Western blot confirmed the absence of NLRC4 protein in the *Nlrc4*^*−/−*^ mice and the presence of protein in the *Nlrc4*^*S533A/S533A*^ and *Nlrc4*^*S533D/S533D*^ lines (Fig.2A). We used FlaTox, a reagent to deliver *Legionella pneumophila* flagellin into the cytosol of cells (Zhao et al., 2011; Rauch et al., 2016; Von Moltke et al., 2012; Ballard et al., 1996), to activate the NLRC4 inflammasome in macrophages. We measured release of the cytosolic enzyme lactate dehydrogenase (LDH) into the supernatant as an assay for pyroptosis downstream of NLRC4-mediated Caspase-1 activation. As expected, we detected increasing levels of LDH release by wildtype C57BL/6J macrophages treated with increasing amounts of FlaTox, while macrophages from our newly generated *Nlrc4*^*−/−*^ mice did not release LDH at any dose of FlaTox (Fig 2B). Macrophages from *Nlrc4*^*S533A/S533A*^ mice showed about a 50% decrease in LDH release at lower concentrations of FlaTox, but did not exhibit a defect in LDH release at the highest FlaTox doses. Macrophages from phosphomimetic *Nlrc4*^*S533D/S533D*^ mice displayed LDH release kinetics comparable to wildtype controls and did not show evidence of spontaneous inflammasome activation in the absence of FlaTox.

**Figure 2.**
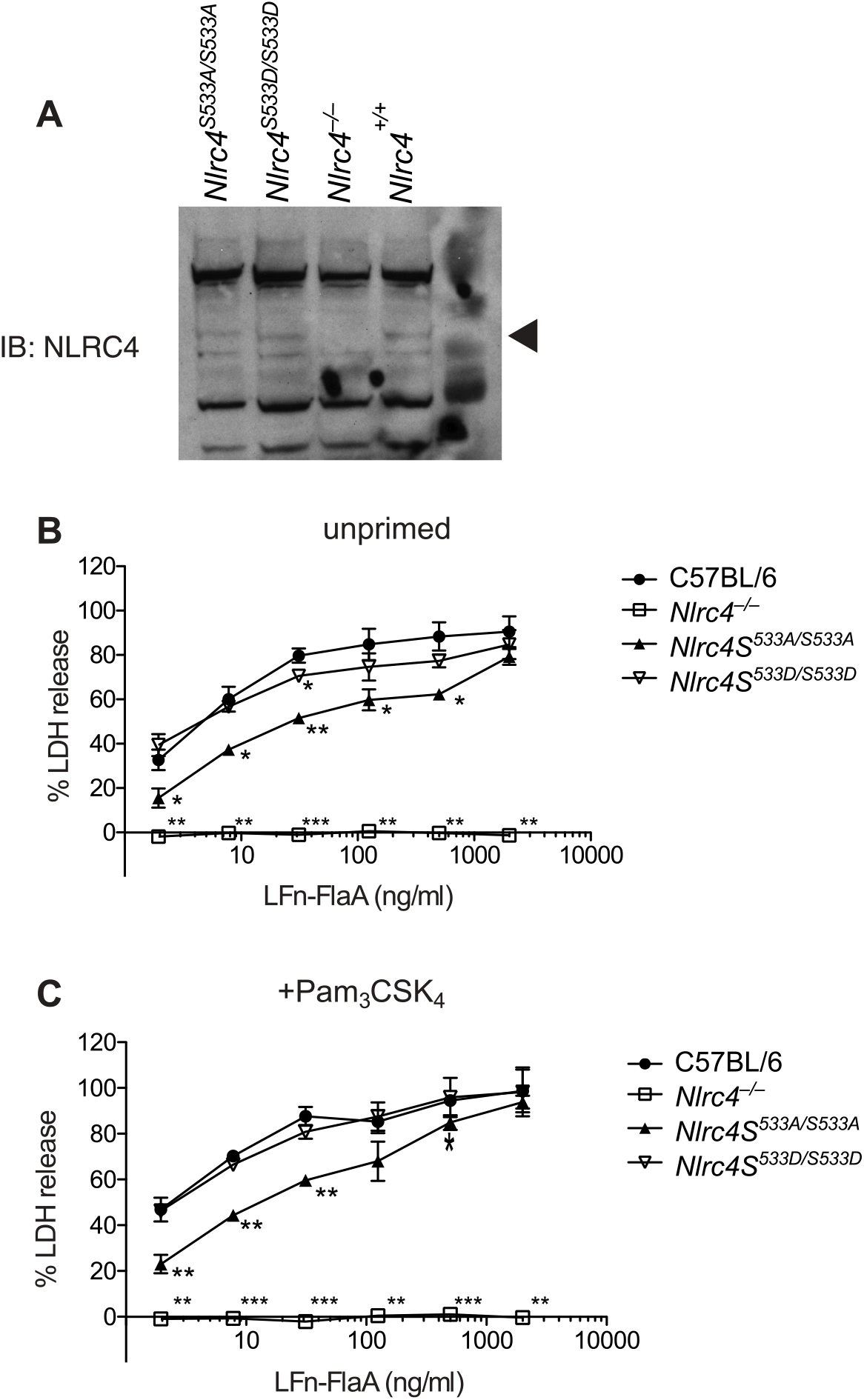
NLRC4 phosphorylation is neither sufficient nor strictly required for response to cytosolic flagellin. (A) Western blot of NLRC4 levels in *Nlrc4* mutant bone marrow-derived macrophages. LDH release into supernatants was measured from macrophages either (B) unprimed or (C) primed 4 hr with 0.5 *μ*g/ml Pam3CSK4 and then treated with 4 *μ*g/ml PA and the indicated dose of LFn-Fla. Data are representative of 3 independent experiments (3 biological replicates per experiment). Mean ± SD. Repeated measures two-way ANOVA vs. C57BL6/J with Dunnett’s multiple comparisons post test, p<0.05, **p<0.005, ***p<0.0005.

We repeated our experiments in macrophages pretreated with the TLR2 ligand Pam3CSK4, as it was suggested that TLR priming may help reveal a function for NLRC4 phosphorylation (Qu et al., 2016). There was no difference between primed and unprimed *Nlrc4*^*S533A/S533A*^ macrophages in LDH release after FlaTox treatment (Fig. 2C). We also did not observe increased pyroptosis in primed *Nlrc4*^*S533D/S533D*^ macrophages compared to controls, either with or without inflammasome stimulation.

We next addressed whether the activation of non-phosphorylatable NLRC4-S533A requires NAIPs, or whether, as previously suggested (Qu et al., 2016), NLRP3 partially compensates for the loss of NLRC4 phosphorylation. We crossed our *Nlrc4*^*S533A/S533A*^ mice to *Nlrp3*^*−/−*^ mice to generate *Nlrc4*^*S533A/S533A*^;*Nlrp3*^*−/−*^ animals. Whether primed or unprimed, *Nlrc4*^*S533A/S533A*^;*Nlrp3*^*−/−*^ macrophages exhibited the same amount of cell death as *Nlrc4*^*S533A/S533A*^ cells upon treatment with a high dose of FlaTox (Fig. 3A). In contrast, *Nlrc4*^*−/−*^ and *Naip1-6*^Δ/Δ^ cells were completely protected, confirming the crucial role of NAIPs in NLRC4 activation (Zhao et al., 2011; Rauch et al., 2016). As a control, treatment of the macrophages with the NLRP3 agonist Nigericin confirmed that we were able to prime NLRP3 activity and that *Nlrp3*^*−/−*^ macrophages lacked NLRP3 activity (Fig. 3B).

**Figure 3.**
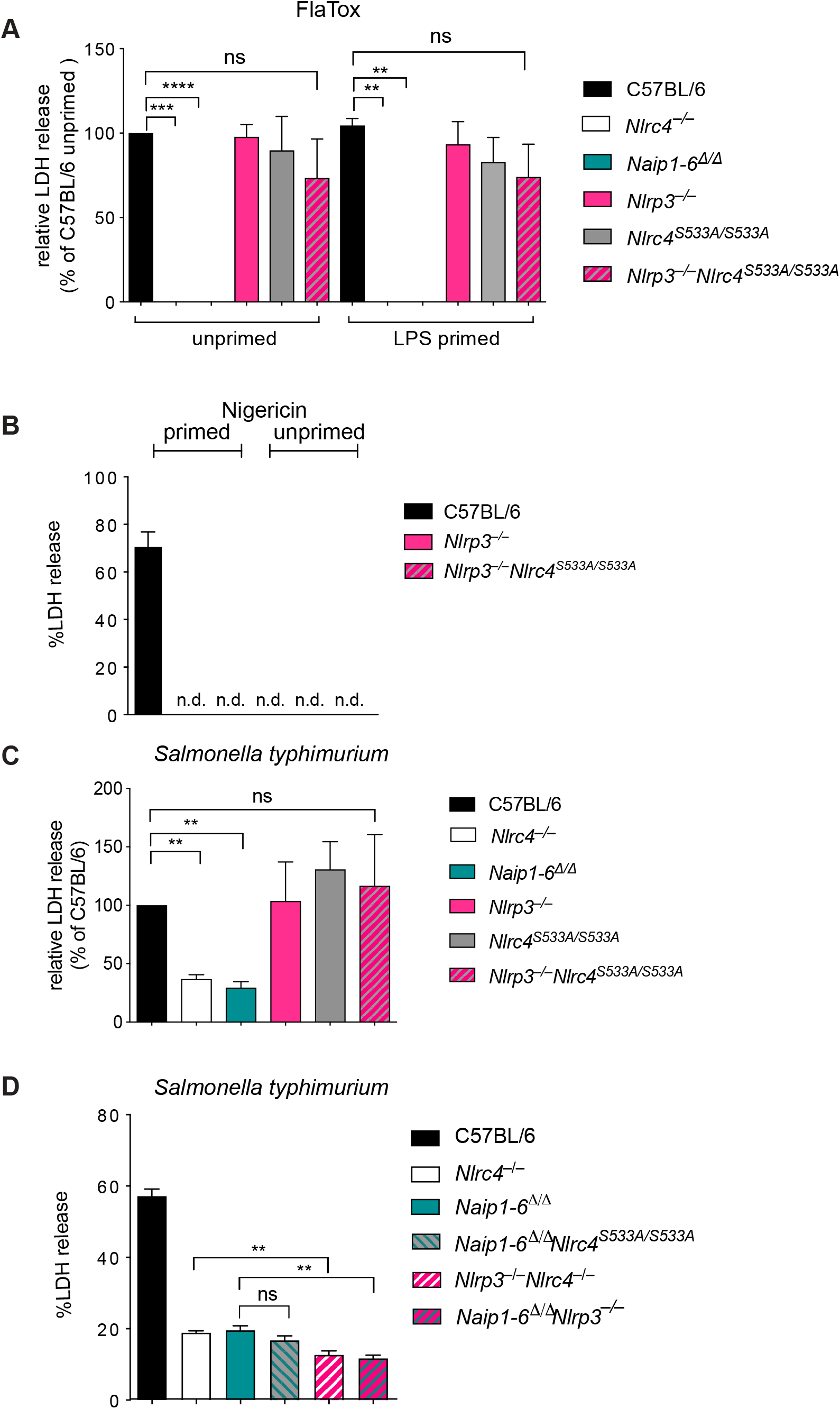
NAIPs, not NLRP3, are required for signaling by NLRC4-S533A. (A) LDH release from macrophages either unprimed or primed for 4 hr with 1 *μ*g/ml LPS and then treated with 2 *μ*g/ml LFn-Fla and 4 *μ*g/ml PA, or (B) with 10*μ*M nigericin, or (C and D) infected with *S. typhimurium* at an MOI of 5 for 4 hr. (A and C) Data are normalized to WT and averaged from 3 independent experiments (3 biological replicates per experiment). Repeated measures one-way ANOVA. (B and D) Data representative of two independent experiments. Mean± SD. Unpaired t-test n.d.: not detectable **p<0.005, ***p<0.0005, ****p<0.00001.

Previous studies of NLRC4 phosphorylation used infection of macrophages with *Salmonella typhimurium* to induce NLRC4 activation, as this pathogen both primes cells via TLR stimuli and activates the NAIP/NLRC4 inflammasome. The use of *Salmonella* as an NLRC4 agonist is complicated because it has previously been shown that *Salmonella* can also activate NLRP3, independently of NLRC4 (Broz et al., 2010). Nevertheless, we also performed *S. typhimurium* infections of our mutant macrophages, closely adhering to the previously published protocol (Qu et al., 2016). However, in contrast to previous results, we failed to detect a significant difference in cell death between *S. typhimurium*-infected *Nlrc4*^*S533A/S533A*^;*Nlrp3*^*−/−*^ and *Nlrc4*^*S533A/S533A*^ macrophages (Fig. 3C). Furthermore, we found no difference between wild-type C57BL6/J and *Nlrc4*^*S533A/S533A*^ macrophages in *S. typhimurium*-induced pyroptosis. As *Salmonella* induced some macrophage cell death even in the absence of NLRC4, we crossed mice to obtain *Nlrc4*^*−/−*^;*Nlrp3*^*−/−*^, *Naip1-6*^Δ/Δ^;*Nlrp3*^*−/−*^ and *Naip1-6*^Δ/Δ^; *Nlrc4*^*S533A/S533A*^ macrophages. Upon infection with *Salmonella*, no major differences in cell death were observed in any of these cells compared to *Nlrc4*^*−/−*^ cells (Fig. 3D). Thus, we conclude there is little if any role for S533 phosphorylation or NLRP3 in NLRC4 activation in macrophages *in vitro*, whereas we confirm that NAIPs are essential for NLRC4 activation.

Macrophage generation *in vitro* relies on conditioned medium that can vary from batch to batch, which may lead to differences in macrophage differentiation. We therefore performed *in vivo* experiments to address the role of NLRC4 phosphorylation. As previously described (Von Moltke et al., 2012; Rauch et al., 2016), wild-type C57BL/6J mice experience acute severe NAIP/NLRC4-dependent hypothermia when injected systemically with FlaTox. This phenotype was not mitigated in *Nlrc4*^*S533A/S533A*^, *Nlrp3*^*−/−*^, or *Nlrc4*^*S533A/S533A*^;*Nlrp3*^*−/−*^ animals, nor was the phenotype exacerbated in *Nlrc4*^*S533D/S533D*^ animals (Fig. 4A). We also performed *in vivo S. typhimurium* infection experiments. Streptomycin-pretreated mice were orally infected with 2*10^7^ bacteria via gavage. As previously reported with an independent *Nlrc4*^*−/−*^ mouse line (Rauch et al., 2017; Sellin et al., 2014), our newly generated *Nlrc4*^*−/−*^ animals show significantly increased numbers of *S. typhimurium* in their cecal tissue and mesenteric lymph nodes, as compared to *Nlrc4*^*+/−*^ littermates (Fig. 4B). We failed to detect any significant difference between *Nlrc4*^*S533A/S533A*^;*Nlrp3*^*−/−*^ and *Nlrc4*^*+/−*^ animals (cohoused) or *Nlrc4*^*S533A/S533A*^;*Nlrp3*^*+/−*^ (littermate) animals. Collectively, the above experiments confirm that our newly generated, C57BL/6J *Nlrc4*-deficient mouse line exhibits no NLRC4 activity in response to *L. pneumophila* flagellin or *S. typhimurium* infection. However, we failed to detect a major role for NLRC4 phosphorylation either during *in vitro* or *in vivo* infection.

**Figure 4.**
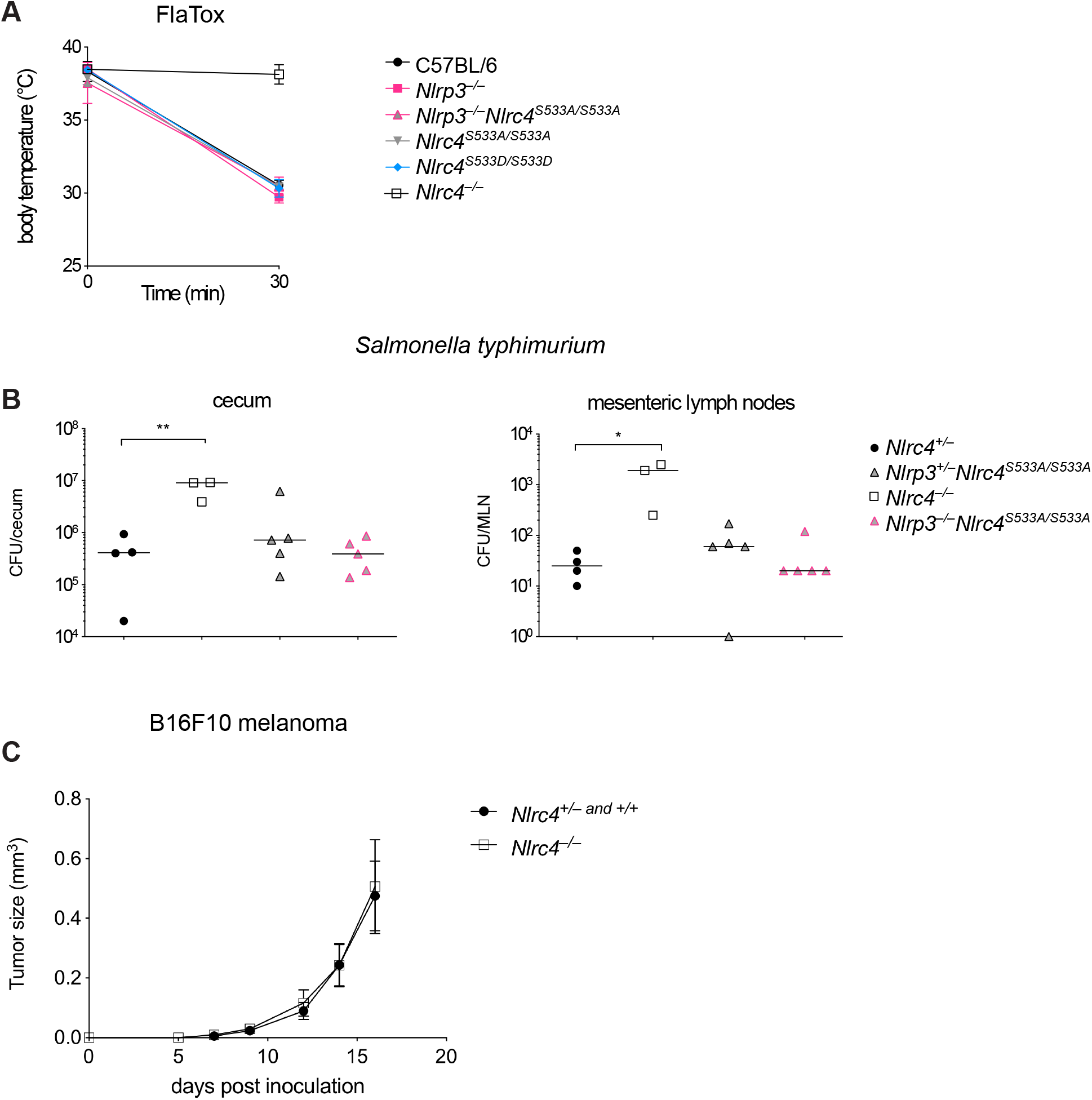
*In vivo* disease susceptibility of *Nlrc4* mutant mice. (A) Rectal temperatures of WT C57BL6/J, *Nlrp3*^*−/−*^, *Nlrc4*^*S533A/ S533A*^;*Nlrp3*^*−/−*^, *Nlrc4*^*S533A/S533A*^, *Nlrc4*^*S533D/S533D*^, and *Nlrc4*^*−/−*^ mice injected retroorbitally with 0.8 *μ*g/g body weight of PA and 0.4 *μ*g/g LFn-Fla (n=3). Mean ± SD are indicated. (B) CFU in cecum and MLN 18 hr after oral *S. typhimurium* infection of *Nlrc4*^*S533A/S533A*^;*Nlrp3*^*−/−*^ and *Nlrc4*^*S533A/S533A*^ littermate mice cohoused with *Nlrc4*^*/–*^ and *Nlrc4*^*−/−*^ littermate mice (n=3-5). Median; Mann-Whitney test, *p < 0.01, **p<0.005. (C) Tumor size of B16F10 melanoma injected subcutaneously into *Nlrc4*-sufficient (*Nlrc4*^+/+^ and *Nlrc4*^*+/−*^) and *Nlrc4*^*−/−*^ littermate mice (n=12). Mean ± SEM. Data are representative of 2 to 3 independent experiments.

Generation of our new *Nlrc4*-deficient mice on a genetic background that is isogenic to C57BL/6J provided the opportunity to address another proposed role for NLRC4: the suppression of cancer progression independently of inflammasome activation. A previous study (Janowski et al., 2016) found that *Nlrc4*^*−/−*^ mice on the C57BL/6N background failed to control subcutaneous growth of the B16F10 melanoma-derived cell line. However, this study did not compare littermates, leaving open the possibility that differences between wild-type C57BL6/J and *Nlrc4*^*−/−*^ mice were due to alterations in the microbiota (Sivan et al., 2015) or other genetic differences between these mice. We therefore challenged our new *Nlrc4*-deficient mice, and their *Nlrc4*-sufficient littermates, subcutaneously with B16F10 melanoma cells and tracked tumor development (Fig. 4C). We did not find any significant differences in tumor size between mice lacking or expressing NLRC4. Thus, we are unable to detect a role for NLRC4 in melanoma suppression.

In conclusion, we do not observe a necessity for NLRC4 phosphorylation or NLRP3 recruitment for inflammasome formation. *Nlrc4*^*S533A/S533A*^ macrophages did show a modest decrease in pyroptosis upon low level ligand stimulation. It is difficult to determine whether this is due to lack of phosphorylation or, alternatively, to mild destablization of the protein due to the S533A mutation. However, since there was no difference observed upon stronger stimulation and the phosphomimetic mutant behaved indistinguishably from wildtype, we conclude that phosphorylation is neither strictly required nor sufficient for NLRC4 activation. We did confirm our previous observation that NAIPs are essential for NLRC4 inflammasome formation (Rauch et al., 2016), which had already been observed in an independently made mouse line (Zhao et al., 2016). We can only speculate on the difference in experimental results with our newly generated *Nlrc4* mutant mice and previous observations. The importance of NLRC4 phosphorylation might only be revealed in a specific genetic context, as the original *Nlrc4*^*−/−*^ mouse line was generated in C57BL/6N embryonic stem cells (Mariathasan et al., 2004). Recently, immunological differences have been described between C57BL/6N and J lines (Simon et al., 2013), which could be explained partially by C57BL/6J carrying a mutation in NLRP12, leading to defects in neutrophil recruitment (Ulland et al., 2016). The original *Nlrc4*^*−/−*^ line might also carry further passenger mutations or have acquired a *de novo* mutation. Such differences might also explain a previous report that *Nlrc4*^*−/−*^ but not *Caspase1*^*−/−*^ fail to control implanted tumors (Janowski et al., 2016), a finding that we could not replicate with our new *Nlrc4*^*−/−*^ mice. Regardless, our results clarify our understanding of NLRC4 by confirming that NAIPs—rather than NLRP3 or phosphorylation—are the main upstream activators leading to inflammasome activation.

## METHODS

### Mice

C57BL/6J (B6) mice were purchased from Jackson laboratories and bred at UC Berkeley. Animal experiments were approved by the UC Berkeley Animal Care and Use Committee. To generate *Nlrc4*-genetically modified mice, fertilized embryos from C56BL/6J mice were injected with Cas9 mRNA and sgRNA, as described (Rauch et al., 2016), along with two DNA oligonucleotides (ssODN) for homology-directed repair. The sgRNA was designed using MIT and Benchling CRISPR design tools and chosen to optimize targeting efficiency relative to efficiency of off-target sites. The following guide was selected: ATTGATTCCTGCCTCCAGAG, non-coding PAM, MIT targeting score = 48, highest off-target score = 5.2. S533 sgRNA was co-injected with S533A (TGCAATGGTTTATCAGCACGGCAGCCTACAAGGACTTTCAGTCACCAAGAGGC CTCTCTGGAGaCAaGAA**<u>gCA</u>**ATtCAGAGTCTGAGAAATACCACTGAGCAAGATGT TCTGAAAGCCATCAATGTAAA TTCCTTC) and S533D (TGCAATGGTTTATCAGCACGGCAGCCTACAAGGACTTTCAGTCACCAAGAGGC CTCTCTGGAGaCAaGAg**<u>gat</u>**ATCCAGAGTCTGAGAAATACCACTGAGCAAGATGT TCTGAAAGCCATCAATGTAAATTCC) ssODNs. ssODNs contained the indicated coding change and several silent point mutations to prevent continued targeting of the repaired allele. ssODNs were synthesized with a terminal 5’ and 3’ phosphorothioate bond for stability and were PAGE purified. Founder mice were genotyped by PCR amplification of *Nlrc4* exon 4 using Ipaf-GenoF (ATGGGTCCAGCATGAACGAG) and Ipaf-GenoR primers (TCTGAGAACAAAT TGATGCCACAC). PCR products were digested with fast alkaline phosphatase (Thermo Fisher) and exonuclease I (NEB) and sequenced with Ipaf-GenoF or Ipaf-GenoR primers. Founders were backcrossed at least three generations to C56BL/6J mice and then crossed to homozygosity.

### Toxins

His-tagged *Bacillus anthracis* protective antigen (PA) and the N-terminus of lethal factor (LF) fused to *L. pneumophila* flagellin (LFn-FlaA) were purified from *Escherichia coli* using Nickel NTA resin, as described previously (Von Moltke et al., 2012). Endotoxin was removed with Detoxi-Gel (Pierce). Toxin doses were 0.8 *μ*g/g body weight of PA combined with 0.4 *μ*g/g LFn-Fla for intravenous (retroorbital) delivery, and as indicated in figure legends for *in vitro* experiments. Rectal temperature was measured at baseline and at 30 min post-injection using a MicroTherma 2T thermometer (Braintree Scientific).

### Tissue culture

Melanoma: The B16.F10 melanoma cells used were from ATCC (CRL-6475). The cells were cultured in DMEM supplemented with 10% fetal bovine serum (FBS), 2 mM glutamine, 100 U/mL penicillin and 100 μg/mL streptomycin, and 1mM sodium pyruvate.

Macrophages: Bone marrow was harvested from mouse femurs, and cells were differentiated into macrophages by culturing in RPMI supplemented with 5% L929 cell supernatant, Glutamine, Pencillin-Streptomycin and 10% FBS in a humidified incubator (37°C, 5% CO2).

### Bacterial strains

*S. typhimurium* strain SL1344 (a gift from G. Barton) was grown overnight in Luria-Bertani broth and was then diluted 1:40 and grown shaking at 37°C to mid-exponential phase (3 hr) to induce expression of the *Salmonella* pathogenicity island 1 T3SS.

### LDH assays

Cytotoxicity was measured via the activity of LDH released from macrophages as described (Lightfield et al., 2008). Macrophages were seeded in 96-well TC-treated plates at 10^5^ cells/well. Where indicated, cells were primed with 0.5 μg/mL Pam3CSK4 or 1μg/ml LPS for 4 hr. For FlaTox treatments, media was replaced with media containing 4 *μ*g/mL PA and the indicated concentrations of LFn-FlaA, cells were incubated 4 hr at 37°C, and supernatants were analyzed for LDH activity. For *Salmonella* infections, *S. typhimurium* were added at an MOI of 5 to a 96-well plate containing 10^5^ macrophages per well. Medium was replaced with medium containing Gentamicin (20 *μ*g/ml) 30 min after infection with *S. typhimurium* to kill extracellular bacteria. Culture supernatants were assayed for LDH activity after 4 hr. For all LDH assays, infection-specific lysis was calculated as the percentage of detergent-lysed macrophages.

### *In vivo Salmonella* infections

*S. typhimurium* infections were done as previously described (Barthel et al., 2003). Briefly, 10- to 15-week-old mice deprived of food and water for 4 hr were gavaged with 25 mg streptomycin sulfate in H_2_O. After 20 hr, mice were again deprived of food and water for 4 hr and were then gavaged with 2× 10^7^ CFUs *S. typhimurium* SL1344. After 18 hr of infection, the animals were sacrificed, tissue was harvested, and ceca were flushed and incubated in PBS containing 400 mg/mL gentamicin for 30 min to kill luminal bacteria before washing 6 times in PBS. Cecum and MLN were homogenized in sterile PBS and plated on MacConkey agar containing 50 *μ*g/mL streptomycin.

### Western blot

Macrophages were grown in tissue culture-treated 24-well plates, media was removed, and cells were lysed in plate in RIPA buffer (50 mM Tris, 150 mM NaCl, 1% NP-40, 0.5% sodium deoxycholate, 0.1% SDS, 1mM PMSF, 1x Roche protease inhibitor tablet [no EDTA], pH 8.0) for 20 min at 4°C. Lysates were clarified by centrifugation at 16,000 × *g* for 10 min at 4°C, and supernatants were separated by 4-12% SDS-PAGE in MES buffer (Invitrogen). Proteins were transferred to Immobilon-FL PVDF membranes (Millipore). Membranes were blocked with 5% milk. Anti-NLRC4 antibody (gift of S. Mariathasan and V. Dixit, Genentech) was detected with a secondary anti-rabbit IgG conjugated to HRP (GE Healthcare).

### Tumor Model

Heterozygous *Nlrc4*^*+/–*^, newly generated on the C57Bl/6J background, were crossed to generate co-housed littermate *Nlrc4*^+/+^, *Nlrc4*^*+/–*^, and *Nlrc4*^*−/−*^ mice. Mice were subcutaneously injected in the right abdominal flank with 1×10^5 B16F10 cells. The tumors were then measured every 2-3 days with digital calipers. Mice were sacrificed 16 to 19 days after injection.

